# Molecular, circuit, and stress response characterization of Ventral Pallidum Npas1-neurons

**DOI:** 10.1101/2021.10.27.466188

**Authors:** Gessynger Morais-Silva, Hyungwoo Nam, Rianne R. Campbell, Mahashweta Basu, Marco Pagliusi, Megan E Fox, Savio Chan, Sergio D Iñiguez, Seth Ament, Marcelo Tadeu Marin, Mary Kay Lobo

## Abstract

Altered activity of the ventral pallidum (VP) underlies disrupted motivation in stress and drug exposure. The VP is a very heterogeneous structure comprised of many neuron types with distinct physiological properties and projections. Neuronal PAS 1-positive (Npas1+) VP neurons are thought to send projections to brain regions critical for motivational behavior. While Npas1+ neurons have been characterized in the globus pallidus external, there is limited information on these neurons in the VP. To address this limitation, we evaluated the projection targets of the VP Npas1+ neurons and performed RNA-seq on ribosome-associated mRNA from VP Npas1+ neurons to determine their molecular identity. Finally, we used a chemogenetic approach to manipulate VP Npas1+ neurons during social defeat stress (SDS) and behavioral tasks related to anxiety and motivation in Npas1-Cre mice. We employed a similar approach in females using the chronic witness defeat stress (CWDS). We identified VP Npas1+ projections to the nucleus accumbens, ventral tegmental area, medial and lateral habenula, lateral hypothalamus, thalamus, medial and lateral septum, and periaqueductal gray area. VP Npas1+ neurons displayed distinct transcriptomes representing distinct biological processes. Chemogenetic activation of VP Npas1+ neurons increased susceptibility to a subthreshold (S)SDS and anxiety-like behavior in the elevated plus maze and open field while the inhibition of VP Npas1+ neurons enhanced resilience to chronic (C)SDS and CWDS. Thus, the activity of VP Npas1+ neurons modulates susceptibility to social stressors and anxiety-like behavior. Our studies provide new information into VP Npas1+ neuron circuitry, molecular identity, and their role in stress response.

## Introduction

The ventral pallidum (VP) is an important structure within the reward circuitry that has a prominent role in the processing of reward information and execution of motivated behaviors (Root et al., 2015; Wulff et al., 2019). From a circuit standpoint, the VP is a structure that is closely connected to the ventral tegmental area (VTA) and the nucleus accumbens (NAc), which are two of the best-characterized areas of the reward circuitry. VP neurons also project to aversive centers such as lateral habenula (LHb). These circuit features make the VP a key structure for encoding hedonic value and control of motivated and aversive behaviors (Faget et al., 2018; Khan et al., 2020). Thus, it is not surprising that altered VP activity is shown to affect both anhedonia and social aversion (Knowland and Lim, 2018). Specifically, pharmacogenetic and optogenetic inhibition of VP parvalbumin-positive neurons (PV+ neurons) projecting to VTA or LHb has been shown to improve depressive-like behaviors in mice (Knowland et al., 2017). In humans, atrophy or lesions in the pallidal regions are related to depression symptoms such as anhedonia and social withdrawal (Miller et al., 2006; Onyewuenyi et al., 2014; Moussawi et al., 2016; Stuke et al., 2016), and VP serotonin binding is reduced in depressed patients (Murrough et al., 2011).

The VP is a highly heterogeneous brain area, comprised of diverse neuronal phenotypes and innervated by numerous brain areas that release many neurotransmitters (Root et al., 2015), intrinsically implicated in its role of controlling complex behaviors. The NAc is the major input source, innervating the VP with GABAergic and peptidergic synapses, which primarily target pallidal GABAergic and, to a lesser extent, cholinergic neurons (Kupchik et al., 2015; Root et al., 2015). The VP neurons also send projections back to the NAc, which are classically known to be GABAergic (Kuo and Chang, 1992; Churchill and Kalivas, 1994). Dopaminergic projections to the VP arise mainly from the VTA (Klitenick et al., 1992; Stout et al., 2016), which also send glutamatergic and GABAergic signals (Taylor et al., 2014; Breton et al., 2019). Both GABAergic and glutamatergic projections from the VP to the VTA are described, while only glutamatergic projections are reported in the VP to LHb connections (Knowland et al., 2017; Tooley et al., 2018).

The Npas1 (Neuronal PAS 1) protein is a transcriptional repressor that has been demonstrated to play an important role in neuronal differentiation (Stanco et al., 2014). It is increased in the brain of rodent embryos and reaches stable levels after the maturation of the nervous system (Teh et al., 2006). In the GPe, the Npas1-positive (Npas1+) neurons are a distinct class of neurons from the more abundant PV+ neurons (Hernandez et al., 2015; Abrahao and Lovinger, 2018; Abecassis et al., 2020) and they strongly project to striatal regions (Glajch et al., 2016). They also receive inputs from diverse brain regions related to stress response, especially from the central amygdala (Hunt et al., 2018). Although the Npas1+ neurons in the VP have not been characterized yet, similarities between the GP and VP can be useful for understanding the properties of these cells in the VP.

Given the pivotal role of the VP in motivated behaviors and its anatomical position, the VP Npas1+ neurons are likely important to the development of behavioral alterations related to stress exposure. Thus, our current work aims to evaluate basic projection patterns and molecular properties in VP Npas1+ neurons, while also characterizing their role in stress and in emotion-related behaviors.

## Material and methods

### Animals

All procedures regarding animal use in this study were approved by the University of Maryland School of Medicine (UMSOM) Institutional Animal Care and Use Committee. Mice were given food and water *ad libitum* and housed in UMSOM animal facilities on a 12:12 h light/dark cycle (experiments performed during the light-phase).

Npas1-Cre-2A-tdTomato hemizygotes (Npas1-Cre) and Npas1-Cre-2A- tdTomato-RiboTag (Npas1-Cre-RT) mice were used as experimental mice. The Npas1- Cre strain was generated as previously described (Hernandez et al., 2015) on a C57BL/6 background. Npas1-Cre-RT mice were generated by breeding Npas1-Cre mice with RiboTag (RT)**^+/+^** mice (Rpl22**^tm1.1Psam/J^**) (Sanz et al., 2009) on a C57BL/6 background. Male CD-1 retired breeders (Charles River, >4 months) were used as aggressors in social defeat stress experiments. Experimental mice were 8 weeks of age at the time of viral injections.

### Drugs

Clozapine-N-oxide (CNO, Cayman Chemical, Ann Arbor, MI, US) was used to activate the designer receptors exclusively activated by designer drugs (DREADDs), diluted in saline (0.1 mg/mL) and injected in a dose of 1 mg/kg (0.1 mL/10 g, i.p.).

### Viral vectors and stereotaxic surgery

Adeno-associated viruses (AAVs) with a double inverted open reading frame containing the excitatory DREADD fused to mCherry hM3D(Gq)-mCherry (AAV5-hSyn-DIO-hM3Dq-mCherry, Addgene plasmid #44362, Watertown, MA, USA) or the inhibitory DREADD fused to mCherry hM4D(Gi)-mCherry (AAV5-hSyn-DIO-hM4Di-mCherry, Addgene plasmid #44361) were used for chemogenetic manipulation of Npas1+ neurons. AAV5-hSyn-DIO-eYFP (UNC Vector Core Facility, Chapel Hill, NC, USA) was used in control groups, and the animals used for tracing of projection targets.

For viral infusions, mice were anesthetized with isoflurane (3% induction, 1.5% maintenance) and had their head fixed in a stereotaxic (Kopf Instruments, Tujunga, CA, USA) for targeting of the VP bilaterally (anteroposterior, +0.9 mm; mediolateral, ±2.2 mm; dorsoventral, -5.3 mm from the Bregma). The virus (300 nL) was infused with 33- gauge Neuros syringes (Hamilton Co., Reno, NV, USA) at a rate of 0.1 µL/min, and the needle was left in place for 12 min following the infusion. Mice were allowed to recover for 2 weeks before the beginning of the experiments.

### Immunohistochemistry

Animals were deeply anesthetized with isoflurane and transcardially perfused with 0.1 M phosphate-buffered saline (PBS) followed by 4% paraformaldehyde (PFA). Brains were removed, post-fixed in PFA for 24 h, and 40 µm sections were collected in PBS using a vibratome (Leica VT1000S, Leica Biosystems Inc., Buffalo Grove, IL, USA). Slices were washed with PBS and blocked in 3% normal donkey serum (NDS) and 0.3% Triton X-100 in PBS for 30 min. After that, slices were incubated in a blocking buffer containing the primary antibodies rabbit anti-mCherry (1:1500; cat. No. 632496, Takara Bio USA, Inc., Mountain View, CA, USA) and chicken anti-GFP (1:1500; #GFP- 1020, Aves Labs Inc., Tigard, OR, USA) at 4 °C overnight. Slices were washed again then incubated in PBS containing the secondary antibodies donkey anti-chicken Alexa 488 (1:1000; cat. No. 703-545-155, Jackson ImmunoResearch Laboratories Inc., West Grove, PA, USA) and donkey anti-rabbit Cy3 (1:1000; cat. No. 711-165-152, Jackson Immuno) at room temperature for 2 hours. Slices were washed then mounted with Vectashield mounting media with DAPI (Vector Laboratories, Inc., Burlingame, CA, USA). Images were captured at 2.5x or 20x magnification on a laser-scanning confocal microscope (Leica SP8, Leica Microsystems, Buffalo Grove, IL, USA).

### Cell-type-specific RNA sequencing

Immunoprecipitated polyribosomes were prepared from the VP of Npas1-Cre-RT and RT mice according to our previous protocol (Chandra et al., 2019; Engeln et al., 2020). RT mice were injected with AAV5-Cre virus into the VP to allow for polyribosome immunoprecipitation. VP punches were collected and pooled from 5 animals. Then the samples were homogenized and 800uμL of the supernatant was incubated in HA-coupled magnetic beads (Invitrogen: 100.03D) overnight at 4u°C. Then the magnetic beads were washed in high salt buffer and RNA was extracted with the RNeasy Micro kit (Qiagen: 74004). For RNA sequencing, samples were checked for RNA quality and only the ones with RNA integrity number > 8 were used. Samples were submitted in biological triplicates for RNA sequencing at the UMSOM Institute for Genome Sciences (IGS) and processed as described previously (Engeln et al., 2020). Libraries were prepared from 10ung of RNA from each sample using the NEBNext Ultra kit (New England BioLabs, Ipswich, MA, USA). Samples were sequenced on an Illumina HiSeq 4000 with a 75ubp paired-end read. Total of 75–110 million reads were obtained for each sample. Reads were aligned to the mouse genome using TopHat (Kim et al., 2013) and then the number of reads that aligned to the predicted coding regions was determined using HTSeq (Anders and Huber, 2010). Significant differential expression was assessed using Limma where genes with the absolute value of log fold change (LFC)u≥u1 and an uncorrected p-value<0.05 in the pairwise comparisons were considered as differentially expressed (Extended Data 1). Transcriptional regulator network was obtained with iRegulon app (Janky et al., 2014) using Cytoscape 3.7.2 software (Shannon et al., 2003), and Gene Ontology functional enrichment analysis using PANTHER Classification System website (Mi et al., 2013) and BiNGO app (Maere et al., 2005) on Cytoscape software (Extended Data 2). Top GO terms were selected based off of highest -log_10_(adjusted-p-value), and Top upstream regulators were selected from the iRegulon generated list if it contained the highest number of predicted of DEGs. To validate enrichment of the NPAS1 gene, we synthetized cDNA using immunoprecipitated samples from Npas1-Cre-RT and RT mice (see above for detailed procedure) using the iScript cDNA synthesis kit (Bio-Rad). cDNA was preamplified using TaqMan Gene Expression Assay system (Applied Biosystems) as previously described (Liu et al., 2014). FAM probe for *npas1* and gadph (20× TaqMan Gene Expression Assay, Applied Biosystems) was used to amplify the products, and then qRT-PCR was run with TaqMan Gene expression Master Mix (2×, Applied Biosystems) on a CFX-384 Touch (Bio-Rad). Quantification of mRNA changes was performed using the ΔΔCT method (Chandra et al., 2015).

### Social defeat stress (SDS)

Social defeat stress was performed as previously by our lab (Chandra et al., 2017; Nam et al., 2019; Fox et al., 2020a, 2020b) using a well-established protocol (Golden et al., 2011). Briefly, experimental mice were placed in hamster cages with perforated clear dividers containing an aggressive resident mouse (retired CD1 breeder). In chronic SDS (CSDS), mice were physically attacked by a resident for 10 min, then housed opposite the resident on the other side of the divider for 24 h sensory interaction. The defeat session was repeated with a novel resident each day for 10 consecutive days. For subthreshold SDS (SSDS) (Chandra et al., 2017; Nam et al., 2019; Fox et al., 2020a) experimental mice were exposed to 3 defeat sessions for 2.5 min each on a single day, separated by 15 min of sensory interaction. 24 hours after the final session, mice were tested for social interaction (SI).

For evaluation of female stress responses, we utilized a vicarious defeat stress paradigm, which is a modified version of SDS to incorporate an emotional stress component to a female mouse by being allowed to witness the defeat of a male counterpart (Iñiguez et al., 2018). In this chronic social witness defeat (CWDS), CSDS is performed as above for 10 min each session, but with a female mouse on the other side of the divider opposite the side where physical attack is taking place. Following the defeat session, the female mouse is moved to a new cage and housed on the other side of the divider opposite from a novel resident, while the male counterpart is housed opposite the resident for sensory interaction as with CSDS. During the entire CWDS, female subjects are only allowed vicarious experience without any physical interactions with male counterparts or the resident CD1s. Defeat session was repeated with a novel resident and a novel female counterpart each day for 10 consecutive days. 24 hours after the final session, female mice were tested for 3-chamber social preference test.

The CSDS, SSDS and CWDS protocols were conduct in a quiet room (<40 dB).

### Behavioral tests

#### Social interaction test (SI)

Mice were placed in an open arena (42 x 42 cm, white walls, and floor) containing a perforated acrylic box (interaction box) centered on one wall. SI was tested first without a social target for 2.5 min, and then for another 2.5 min immediately after with a novel CD1 mouse serving as a social target. Time spent in the interaction zone (virtual area of 9 cm surrounding the interaction box) and the corner zone (corners of 9 cm opposite from the interaction zone) was measured using a video tracking software (TopScan Lite 2.0, CleverSys, Reston, VA, USA).

#### 3-Chamber Social Preference Test

Mice were placed in a rectangular arena (60 x 40 cm, white walls and floor) divided into 3 chambers (20 x 40 cm) by perforated clear acrylic dividers. The two outer chambers contain wire mesh cups, while the central chamber is empty. The experimental mouse is first placed in the central chamber of the arena with two empty wire mesh cups and allowed to explore for 5 min. Then the experimental mouse is allowed to explore the arena for an additional 5 min, this time with unfamiliar sex- and age-matched mouse in one of the wire mesh cups. The amount of time spent in the chamber containing the cups (empty or novel mouse) is measured using a video tracking software (TopScan Lite).

#### Forced swim test (FST)

Mice were placed in a glass cylinder (20 cm diameter) ¾ filled with water at 25±1 °C and recorded for 6 min. Time spent immobile was scored by a blinded experimenter using the software X-Plo-Rat 2005 (developed by the research team of Dr. Morato, FCLRP, USP, Brazil).

#### Splash test (ST)

Mice were placed in an empty glass cylinder (20 cm diameter) and had their dorsal coat sprayed three times with a viscous sucrose solution (10% sucrose v/v). Behavior was recorded for 5 min to the evaluation of time spent grooming by a blinded experimenter using the software X-Plo-Rat 2005.

#### Elevated plus maze (EPM)

The EPM apparatus is a plus-shaped acrylic apparatus, elevated 50 cm above floor level with two open arms (30 cm length x 5 cm width) and two opaque closed arms (30 cm length x 5 cm width x 15 cm height) connected by a common central platform (5 cm length x 5 cm width). Mice were placed in the center of the apparatus, facing an open arm, and their behavior was recorded over 5 min to the evaluation of spatiotemporal measures, i.e., number of entries (arm entry = four paws in the arm) into the closed arms, percentage of entries into the open arms, percentage of time spent in the open arms and risk assessment behaviors, i.e., frequency of head dips and stretch attend posture (Morais-Silva et al., 2016). Behavioral analyses were performed by a blinded experimenter using the software X-Plo-Rat 2005.

#### Open field (OF)

To evaluate the locomotor activity and anxiety-like behavior, mice were put in the same open arena used for SI (missing the interaction box), virtually divided into a peripheric and a central zone (center square, 15 x 15 cm), for 5 min for the time spent and the distance traveled in each zone. Behavioral analyses were performed using TopScan Lite.

#### Sucrose preference test (SPT)

This protocol lasted for 3 days and was performed as follows: on the first day, mice were given two conical 50 mL tubes with sipper tops containing water for 24 hours. The next day, bottles were substituted with one filled with a 1% sucrose solution and one with water for 48 hours. Bottles were rotated daily to account for side preference. Sucrose preference (amount of sucrose solution consumed over water) was determined by weighing the tubes right before offering to the animals and after the end of the experiment. Solution lost by leakage or evaporation was determined by a tube placed in an empty box at the same time as the solution was offered to the animals.

#### Statistical analyses

Data were expressed as mean±s.e.m and statistical analysis was performed using Statistica 7 (StatSoft Inc., Tulsa, OK, USA) and GraphPad Prism 6. Depending on the experiment, data were analyzed by two-way ANOVA considering the factors virus (eYFP or DREADDs) and defeat (control or defeat), by the repeated-measures ANOVA considering the repeated factor session (no target or target) or by the Student’s t-test. The test used is identified in the results section for each experiment. In cases where ANOVA showed significant differences (p ≤ 0.05), the Newman-Keuls *post-hoc* test was performed. Samples were excluded if animals had an inappropriate viral placement or if they failed Grubbs’ outlier test. Sample sizes were determined from previous studies (Nam et al., 2019; Fox et al., 2020a).

## Results

### Viral tracing of ventral pallidum Npas1+ neuron projection targets

To identify projection targets of Npas1+ neurons we infused AAV-DIO-eYFP into the VP of Npas1-Cre mice, which contain TdTomato fluorophores within Npas1+ cells. After confirming the injection site in the VP (Figure 1A, also available in low magnification in Figure 1B), we were able to identify eYFP-positive neuronal fibers multiple brain regions. eYFP-positive neuronal fibers were observed within the NAc, medial and lateral septum, medial and lateral habenula, lateral hypothalamus, thalamic nuclei (mediodorsal, paraventricular and centrolateral), VTA, periaqueductal gray area, and within the medial anterior olfactory nucleus (AOM) and lateral anterior olfactory nucleus (AOL) (Figure 2). Among these regions, fibers were found to be close to Npas1+ neurons within the NAc, the lateral hypothalamus, anterior olfactory nucleus, and the lateral septum. Other regions did not contain many Npas1+ cells.

**Figure 1.**
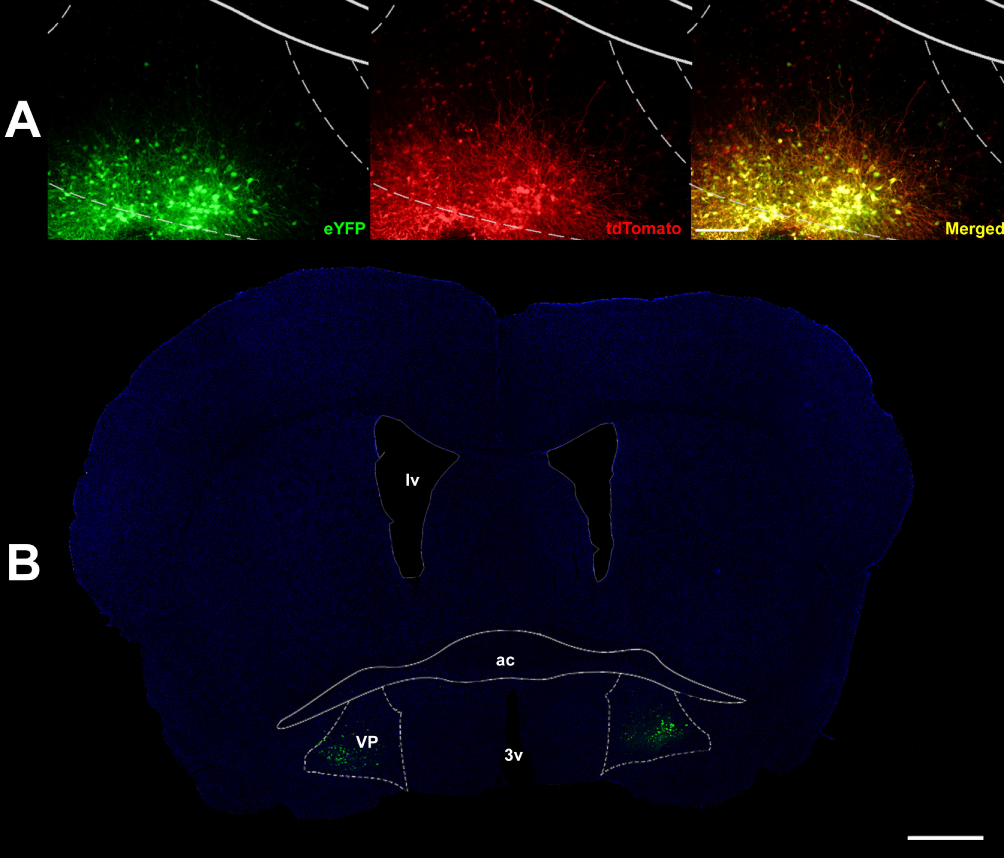
Representative images of the injection site in the ventral pallidum (VP). The first column of panel A (from the left) shows the expression of the eYFP (in green), while the second column shows the expression of the tdTomato in Npas1 neurons (in red). The third column shows the merged images. Panel B is representative of the injection site in the VP at low-magnification (2.5x). Scale bar represents 1mm. ac, anterior commissure. lv, lateral ventricle. 3v, third ventricle. VP, ventral pallidum.

**Figure 2.**
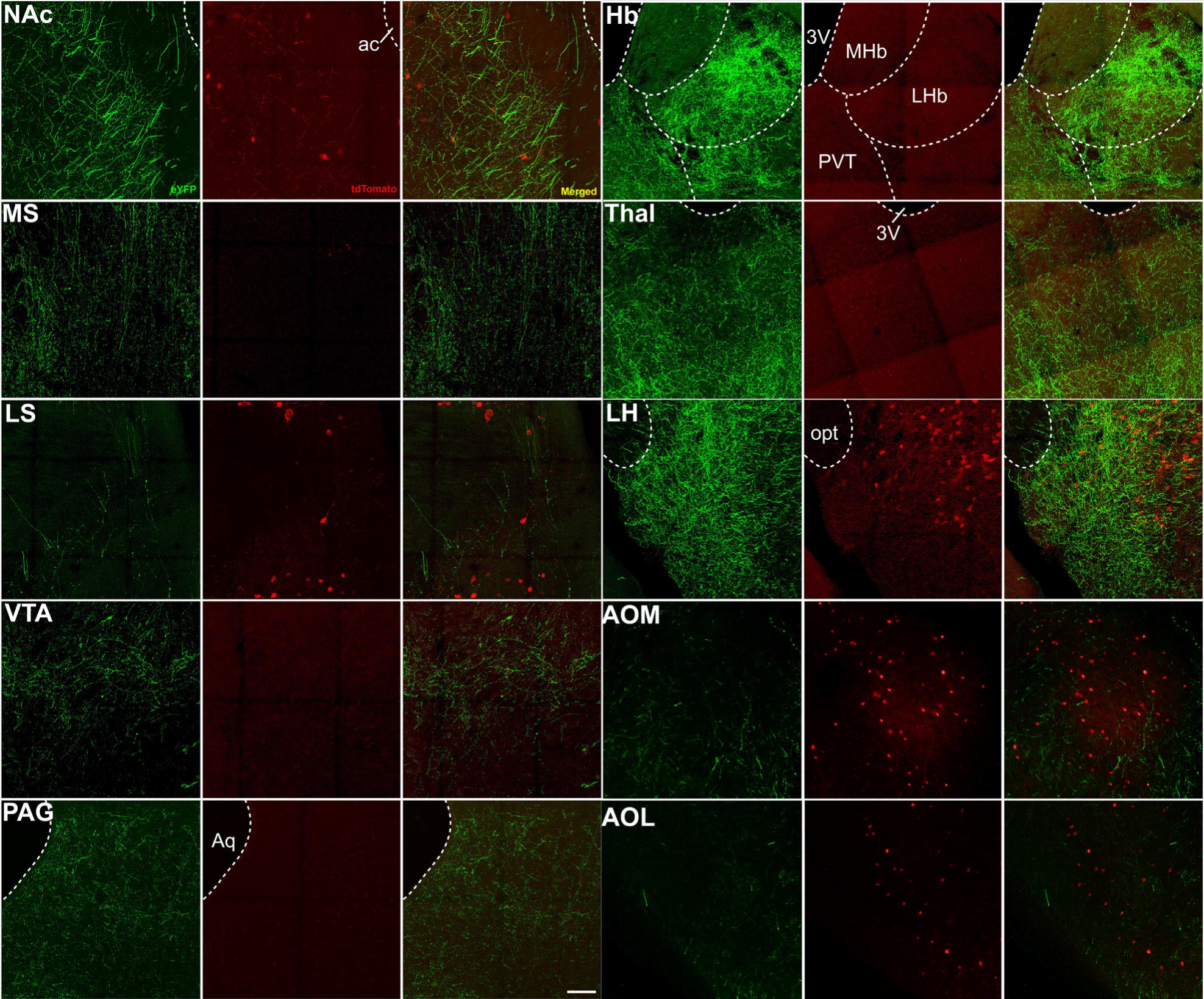
Representative images of DIO-eYFP and tdTomato expression in multiple brain regions of Npas1-cre mouse. The first and fourth columns (from the left) show the expression of the eYFP (in green) within the brain regions, while the second and fifth columns show the expression of the tdTomato (in red). The third and sixth columns show the merged images. Scale bar represents 100 µm. NAc, nucleus accumbens. ac, anterior commissure. LS, lateral septum. MS, medial septum. Hb, habenula. MHb, medial habenula. LHb, lateral habenula. PVT, paraventricular nucleus of the thalamus. LH, lateral hypothalamus. opt, optical nerve. Thal, thalamus. LV, lateral ventricle. 3V, third ventricle. VTA, ventral tegmental area. PAG, periaqueductal gray. Aq, cerebral aqueduct. AOM, anterior olfactory nucleus, medial part. AOL, anterior olfactory nucleus, lateral part.

### Genetic profiling of ventral pallidum Npas1+ neurons

Using Npas1-Cre-RiboTag (RT) mice, we isolated ribosome-associated mRNA from VP Npas1+ neurons, which was analyzed for their genetic profile relative to all VP neurons. This approach allows for cell-type mRNA enrichment (Sanz et al., 2009; Chandra et al., 2015), and allowed us to specifically analyze the genetic profile of the Npas1+ neurons. mRNA used for sequencing was used for validation of Npas1 gene enrichment, as it was confirmed that there was a 4.98-fold enrichment (Figure 3A, t_8_ = 7.881, p < 0.0001). Out of 23,843 total genes detected, 2,297 genes were identified as differentially expressed (DEGs, Figure 3B), of which 562 genes were identified as significantly enriched in Npas1+ neurons compared to nonspecific VP neurons (Extended Data 1). Top entries from Biological Process GO term analysis based on p-value were nervous system development and neurogenesis, and the top terms from Molecular Function section were protein binding, cytoskeletal protein binding, and chloride channel activity (Figure 3C, Extended Data 2). To further investigate whether these enriched genes are regulated by a common transcription factor in Npas1+ neurons within the VP, upstream regulator prediction analysis was performed. Transcription factors SOX6 and SMAD2 had the highest number of predicted DEG targets (Figure 3D). Several SOX6 and SMAD2 targets, including *Sema6a* and *Sox6* were associated with the Top GO terms, Neurogenesis and Development of Nervous System.

**Figure 3.**
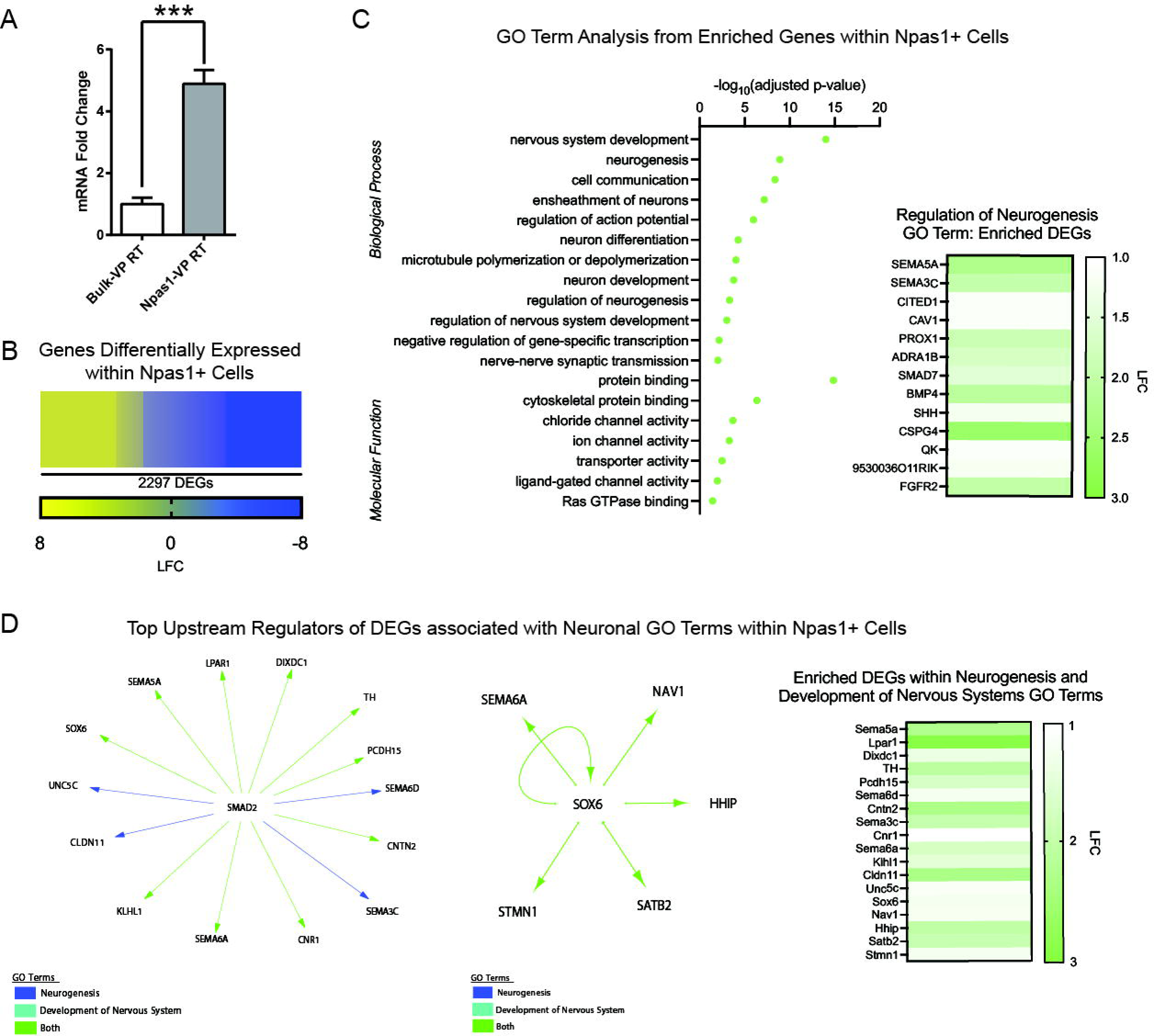
Cell-type specific profiling of mRNA from VP Npas1+ neurons reveal potential functions and regulators these neurons. **A.** Validation of Npas1 enrichment in ribosome-associated mRNA from VP Npas1+ neurons used in genetic profiling. **B.** Overview of all differentially expressed genes (DEGs) identified with the distribution of log fold change (LFC) within DEGs. **C.** Top hits from Biological Process and Molecular Function GO term analysis and an example from one of the top hits, regulation of neurogenesis, with the degree of fold change of genes within the GO term. **D.** 2 Top hits from upstream regulator prediction analysis using top 2 GO terms identified, neurogenesis and development of nervous system. Predicted DEG targets of each regulator are identified with arrows along with their fold change analyzed in VP Npas1+ neurons.

### Bidirectional Ventral pallidum Npas1+ neuron chemogenetic manipulation during subthreshold social defeat stress

To evaluate the role of VP Npas1+ neurons in stress behavior, Npas1-Cre male mice received the excitatory DREADD hM3Dq into the VP and, 30 min before the SSDS, received 1 mg/kg of CNO *i.p*. On the following days, social interaction (SI) was measured followed by the forced swim test (FST) (Figure 4A). Chemogenetic activation of VP Npas1+ neurons induced social avoidance in SSDS mice compared to non-stressed mice. Repeated measures ANOVA revealed a significant effect of stress (F_1,30_=4.42, p<0.05), interaction between virus and stress (F_1,30_=10.68, p<0.01), and an interaction between virus, stress and SI session (F_1,30_=4.25, p<0.05) for time spent in the interaction zone (Figure 4B). Using *post hoc* comparisons, the hM3Dq-SSDS group showed a decrease in the time spent in the interaction zone when the target was present compared to the eYFP-Control, eYFP-SSDS, and hM3Dq-Control. Repeated measures ANOVA revealed a significant effect of the interaction between virus and stress (F_1,30_=9.66, p<0.01), the SI session (F_1,30_=10.20, p<0.01) and the interaction between virus, stress, and SI session (F_1,30_=6.33, p<0.05) for the time spent in the corner zone (Figure 4C). The hM3Dq-SSDS group showed an increase in time spent in the corner zone when target was present compared to eYFP-Control, eYFP-SSDS, hM3Dq-Control, and compared to itself when the target was absent.

**Figure 4.**
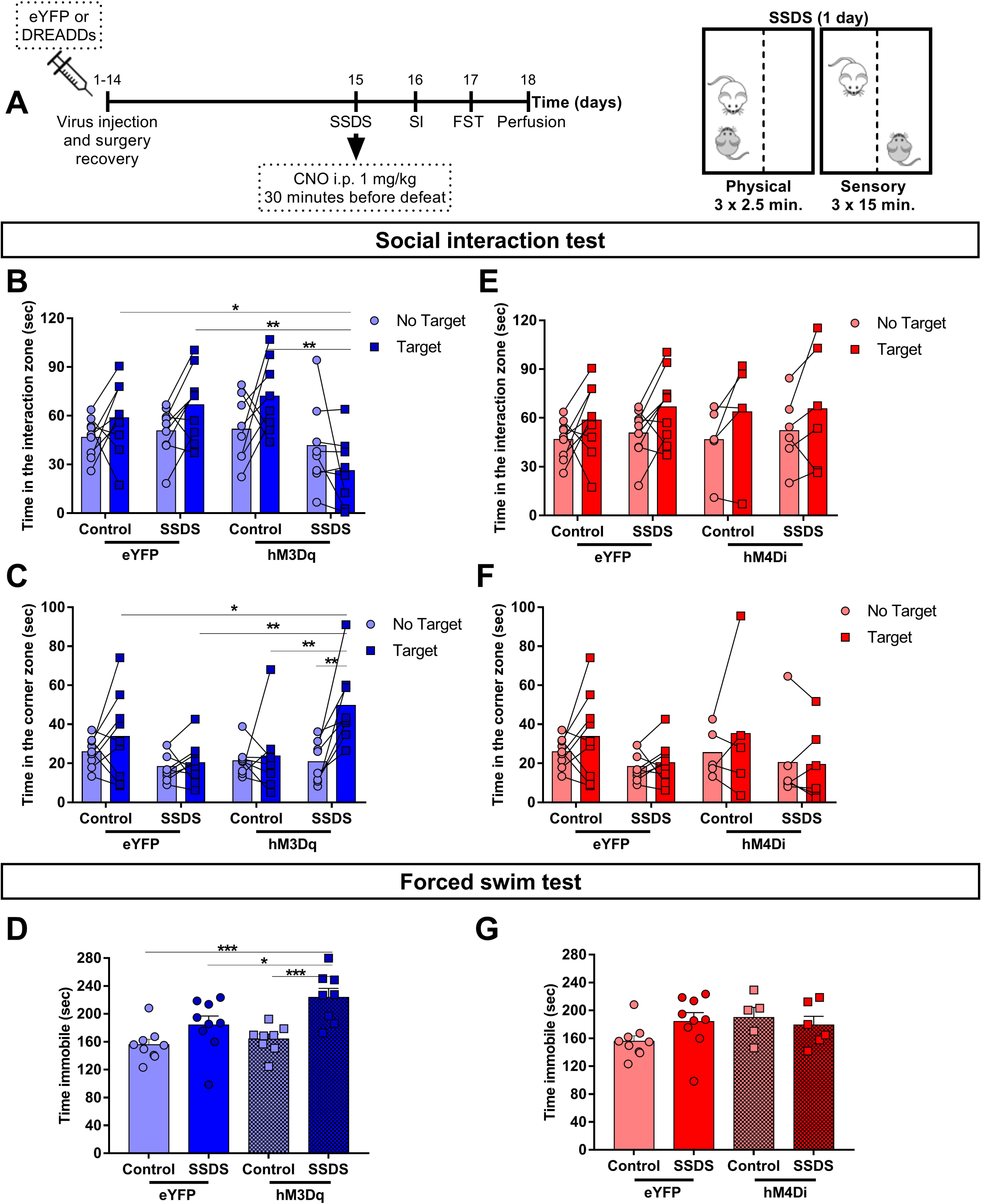
Chemogenetic activation of VP Npas1 neurons increases susceptibility to the subthreshold social defeat stress (SSDS) in male mice. **A**, representative timeline of the experiments involving the manipulation of VP Npas1 neurons during the SSDS. **B, C, D**, effects of the chemogenetic activation of VP Npas1 neurons using the hM3Dq DREADDs during the SSDS in the social interaction (SI) and forced swim (FST) tests. **E**, **F, G,** effects of chemogenetic inhibition of the VP Npas1 neurons using the hM4Di DREAADs during the SSDS in the SI and FST. Data is presented as mean + the individual values obtained for each animal (n = 5-9 animals per group). *, **, *** indicates p < 0.05, p < 0.01, p < 0.001, respectively.

In the FST, chemogenetic activation of VP Npas1+ neurons induced an increase in time immobile in SSDS animals. Two-way ANOVA revealed a significant effect for virus (F_1,30_=5.08, p<0.05) and stress (F_1,30_=17.41, p<0.001) in time spent immobile during the FST (Figure 4D). The *post hoc* test showed an increase in time spent immobile in the hM3Dq-SSDS group compared to eYFP-Control, eYFP-SSDS, and hM3Dq-Control.

We further evaluated chemogenetic inhibition of VP Npas1+ neurons during SSDS. Npas1-Cre male mice received the inhibitory DREADD hM4Di into the VP and, 30 min before the SSDS, received 1 mg/kg of CNO *i.p*. Chemogenetic inhibition of the VP Npas1+ neurons did not alter the behavioral response to SSDS in both the SI and FST. Repeated-measures ANOVA showed a significant effect of the SI session (F_1,26_=12.91, p<0.001) for time spent in the interaction zone (Figure 4E), indicating an increase in the time spent in the interaction zone when the target was present in all groups. No significant alterations were found for time spent in the corner zone (p>0.1) (Figure 4F) and for time spent immobile during the FST (Figure 4G).

### Ventral pallidum Npas1+ neuron chemogenetic inhibition during chronic social defeat stress

Since chemogenetic activation of VP Npas1+ neurons induced a stress susceptible outcome after a subthreshold stress, we next examine if inhibiting these neurons during repeated stress, CSDS, could prevent stress susceptible behavior. Npas1-Cre male mice received the inhibitory DREADD hM4Di into the VP and received 1 mg/kg of CNO *i.p*. 30 min before each defeat session during 10 days of CSDS. On the day after the final defeat session of CSDS, SI was performed followed by FST 24 hours later (Figure 5A). VP Npas1+ neuron chemogenetic inhibition blocked stress-induced social avoidance in CSDS mice. Repeated-measures ANOVA showed a significant effect of the interaction between virus and SI session (F_1,23_=4.85, p<0.05), stress and SI session (F_1,23_=10.89, p < 0.01) and virus, stress, and SI session (F_1,23_=4.22, p=0.05) for the time spent in the interaction zone (Figure 5B). The eYFP-CSDS group spent less time in the interaction zone when the target was present compared to the eYFP-Control, hM4Di-Control, hM4Di-CSDS, and, compared to itself when the target was absent. No significant alterations were found for time spent in the corner zone (p>0.1) (Figure 5C).

**Figure 5.**
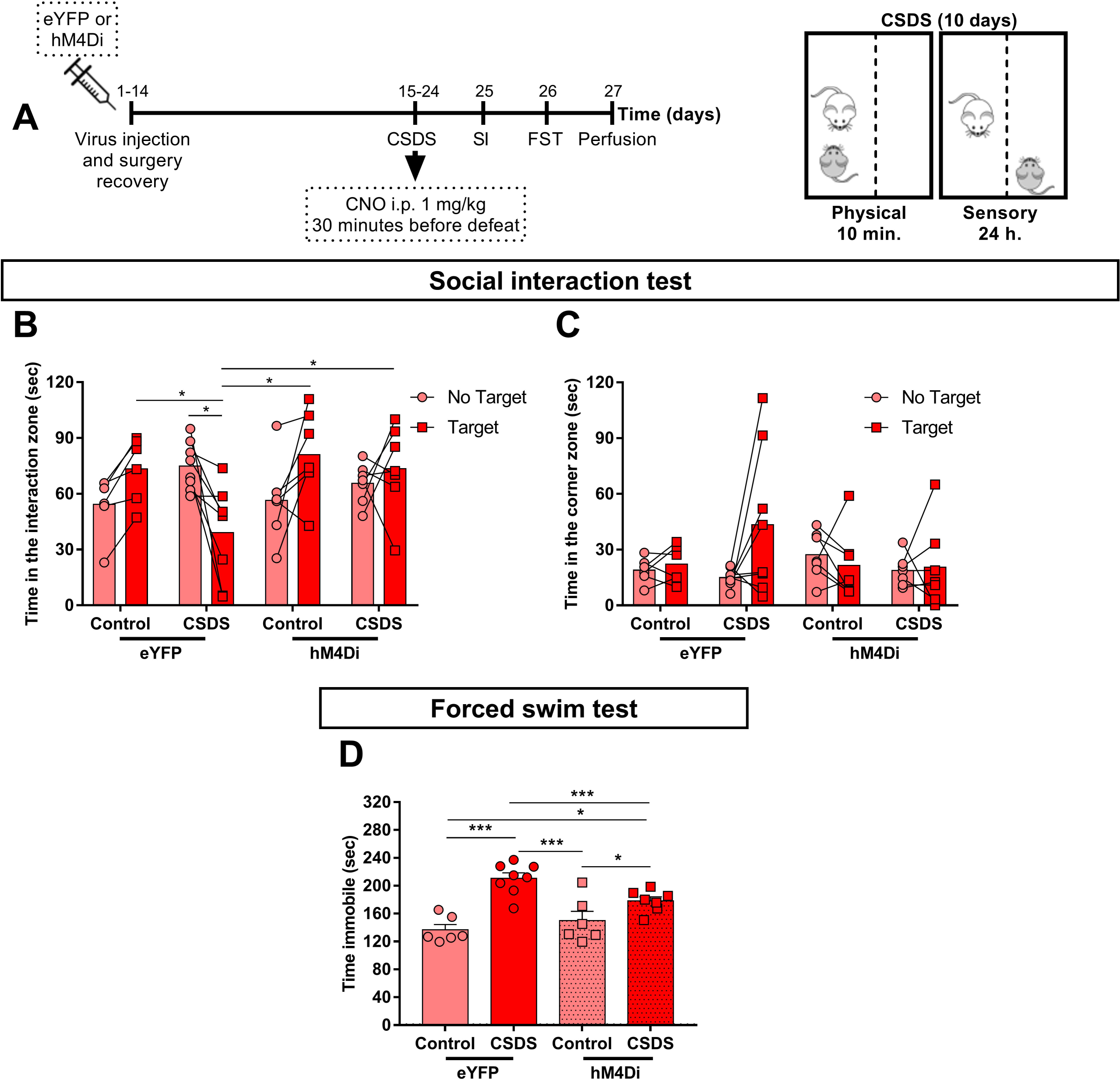
Chemogenetic inhibition of VP Npas1 neurons decreases susceptibility to the chronic social defeat stress (CSDS) in male mice. **A**, representative timeline of the experiments involving the inhibition of the VP Npas1 neurons during the CSDS. **B, C, D,** effects of the chemogenetic inhibition of VP Npas1 neurons using the hM4Di DREAADs during the CSDS in the SI and FST. Data is presented as mean + the individual values obtained for each animal (n = 6-8 animals per group). *, **, *** indicates p < 0.05, p < 0.01, p < 0.001, respectively.

Chemogenetic inhibition of VP Npas1+ neurons reduced time immobile in the FST after CSDS (Figure 5D). Two-way ANOVA showed a significant effect for stress (F_1,23_=33.26, p<0.001) and interaction between virus and stress (F_1,23_=6.65, p<0.05). The eYFP-CSDS group spent more time immobile when compared to the eYFP-Control, hM4Di-Control, and hM4Di-CSDS groups, while the hM4Di-CSDS group spent more time immobile compared to the eYFP-Control and hM4Di-Control groups.

### Ventral pallidum Npas1+ neuron chemogenetic activation during behavioral tests related to anxiety and motivation

To characterize baseline behavioral effects of VP Npas1+ neuron activation, Npas1-Cre male mice received the excitatory DREADD hM3Dq into the VP and 30 min before each behavioral test, received 1 mg/kg of CNO *i.p* (Figure 6A). In the EPM, the number of closed arm entries was not affected by the chemogenetic activation (t_16_=0.35, p>0.5) (Figure 6B), as well as the percentage of entries (t_16_=1.29, p>0.1) (Figure 6C, left) and time (t_16_=0.77, p>0.1) (Figure 6C, right) in the open arms. However, activation of the VP Npas1+ neurons increased the frequency of stretch attend postures (t_16_=2.31, p<0.05) (Figure 6D, left), decreased the frequency of head dips (t_16_=2.29, p<0.05) (Figure 6D, right) and increased the percentage of protected stretch attend postures (t_16_=2.08, p=0.05) (Figure 6E) in the EPM. In the OF, chemogenetic activation of VP Npas1+ decreased the distance travelled in the center (t_16_=2.24, p<0.05) (Figure 6F, left) but not the total distance travelled (t_16_=1.61, p>0.1) (Figure 6F, right) nor the time spent in the center (t_16_=0.67, p>0.5) (Figure 6G). Social interaction was not affected by VP Npas1+ neuron activation, similar to non-stressed animals in Figure 4. There was a significant effect of SI session for the time spent in the interaction zone (F_1,16_=15.78, p<0.01) (Figure 6H) and for the time spent in the corners (F_1,16_=6.99, p<0.01) (Figure 6I), indicating an increase in time spent in the interaction zone when the target was present in all groups. There was no significant alteration in time spent grooming in the ST (t_16_=0.97, p>0.1) (Figure 6J), sucrose preference (t_16_=0.13, p>0.5) (Figure 6K) or sucrose consumption in the SPT (t_16_=0.43, p>0.5) (Figure 6L).

**Figure 6.**
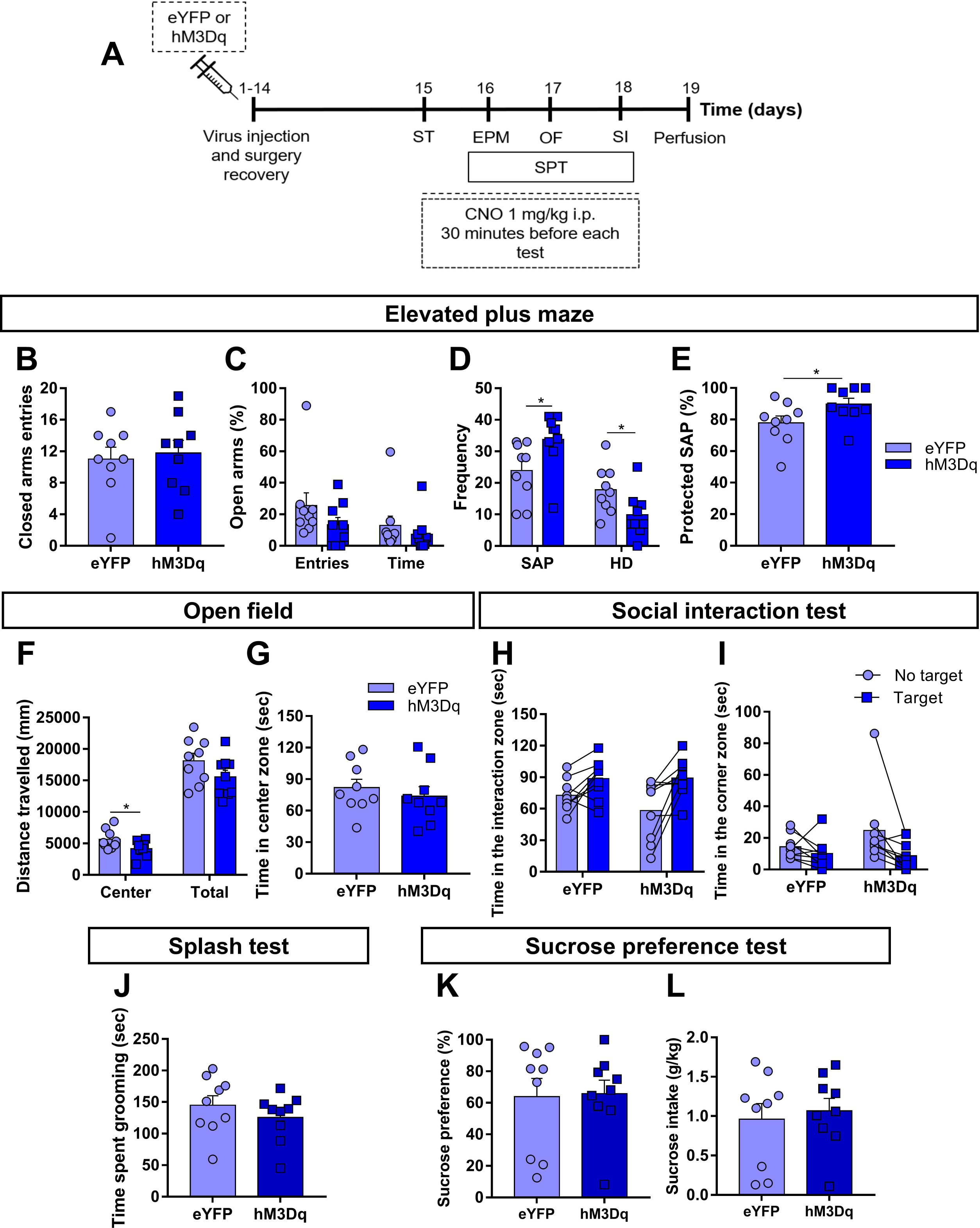
Chemogenetic inhibition of VP Npas1 neurons is anxiogenic in male mice. **A**, representative timeline of the experiments involving the activation of VP Npas1 neurons during behavioral tests that evaluate anxiety-like and depressive-like behaviors. **B, C, D, E,** effects of chemogenetic inhibition of VP Npas1 neurons using the hM4Di DREAADs during the elevated plus maze (EPM) test. **F, G,** effects of the chemogenetic inhibition of the VP Npas1 neurons using the hM4Di DREAADs during the open field (OF) test. **H, I,** effects of the chemogenetic inhibition of VP Npas1 neurons using the hM4Di DREAADs during the SI. **J,** effects of chemogenetic inhibition of VP Npas1 neurons using the hM4Di DREAADs during the Splash test (ST). **K, L,** effects of chemogenetic inhibition of VP Npas1 neurons using the hM4Di DREAADs on the sucrose preference and intake on the sucrose preference test (SPT). Data is presented as mean + the individual values obtained for each animal (n = 8-9 animals per group). * indicates p < 0.05.

### Ventral pallidum Npas1+ neuron chemogenetic inhibition during the chronic witness defeat stress in females

To examine the effects of chemogenetic inhibition of VP Npas1+ neurons during chronic social stress in females, female Npas1-Cre mice received the inhibitory DREADD hM4Di into the VP and received 1 mg/kg of CNO *i.p*. 30 min before each witness defeat session during 10 days of CWDS. On the day after the final defeat session of CSDS, social preference was tested in the 3-chamber social preference test (Figure 7A). In this test, female mice that did not receive inhibitory DREADD (eYFP) reduced their social interaction after they underwent CWDS (Figure 7B). However, mice that received inhibitory DREADD hM4Di into the VP were not affected by CWDS and showed similar levels of social preference compared to the non-stressed control mice.

**Figure 7.**
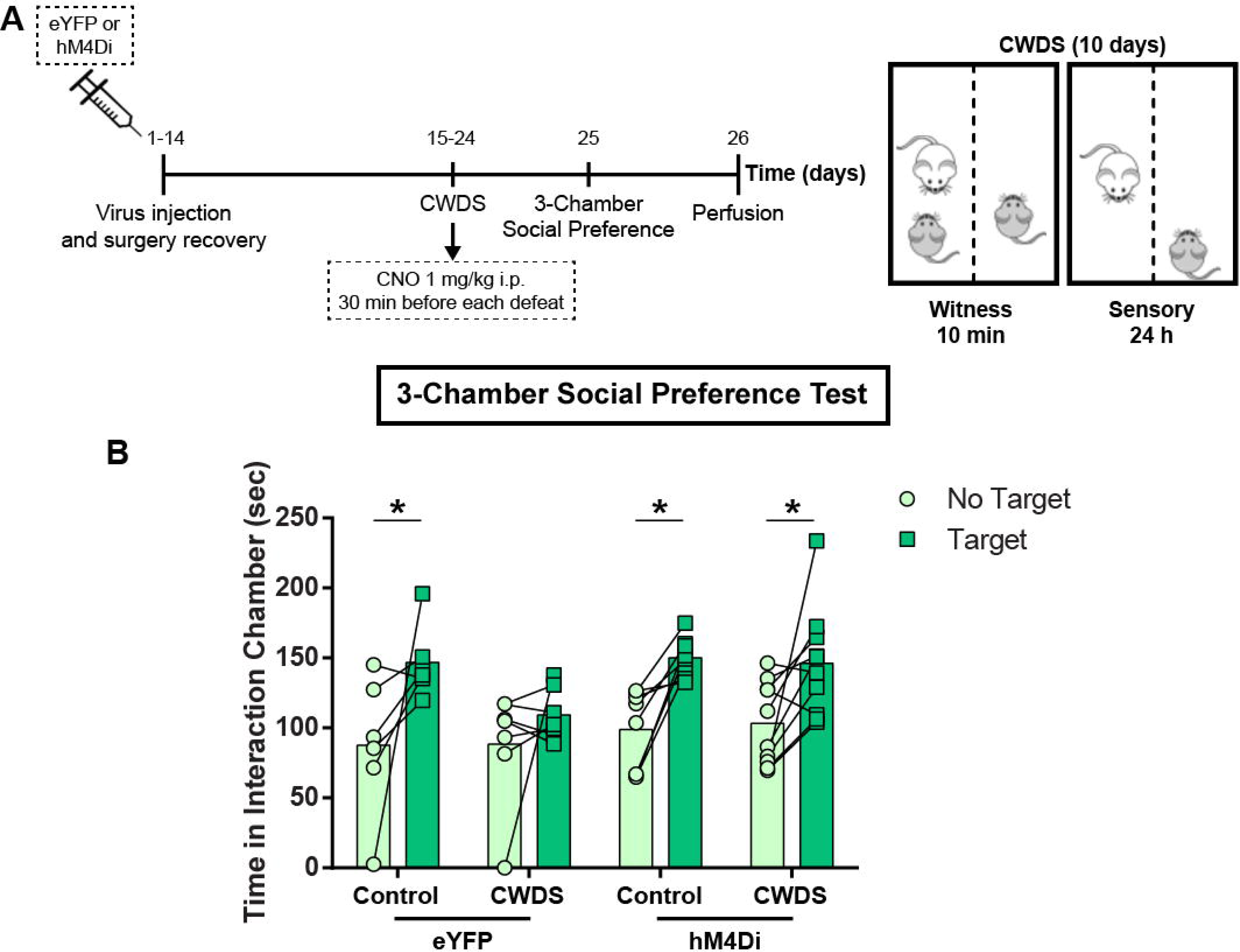
Chemogenetic inhibition of VP Npas1 neurons decreases the susceptibility to chronic witness defeat stress (CWDS) in female mice. **A**, representative timeline of the experiments involving the inhibition of VP Npas1 neurons during the CWDS. **B,** effects of the chemogenetic inhibition of the VP Npas1 neurons using the hM4Di DREAADs during the CWDS in the 3-chamber social preference test. Data is presented as mean + the individual values obtained for each animal (n = 6-10 animals per group). *, p < 0.05.

All the groups except for the eYFP-CWDS group increased their interaction when the target animals were present (Figure 7B). The repeated-measures ANOVA showed a significant effect of the session (F_1,27_=21.77, p<0.0001) but no significant effects of virus treatment or interaction (F_3,27_=2.76 p=0.061 and F_3,27_=0.71, p=0.55, respectively). *Post hoc* test showed significantly increased interaction time with the target present in all groups except for the eYFP-CWDS group.

## Discussion

In our study we identified distinct connections of the VP Npas1+ neurons. While little information is known about the projection targets and cellular identity of these VP neuron subtypes, similarities between the GPe and the VP can aid our understanding of the cellular proprieties of the VP Npas1+ neurons. Both regions originate from similar progenitors and are comprised of glutamatergic, cholinergic, and GABAergic neurons, the latter comprising the majority of VP neurons (Root et al., 2015; Ma and Geyer, 2018). In the GPe, GABAergic neurons are classified into three subgroups, highly segregated according to their electrical properties and the expression of genetic markers (Abrahao and Lovinger, 2018). Among these markers, the Npas1 and PV markers differentiate into two projection patterns. The Npas1+ neurons constitute the principal projection to the dorsal striatum while the PV+ neurons project mainly to the subthalamic nucleus (Hernandez et al., 2015; Glajch et al., 2016). There are also projections of the GPe Npas1+ neurons to the somatosensory, somatomotor, and orbital cortex (Abecassis et al., 2020). Our results revealed that, compared to the GPe, Npas1+ neurons in the VP have a broader and more ventral pattern of projection, without projecting to cortical areas. Others have found similar neuronal projections when examining the entire VP (Knowland et al., 2017; Faget et al., 2018; Wulff et al., 2019). This connectivity suggests that these neurons in the VP may regulate multiple facets of emotional processing through their various outputs.

We additionally analyzed the cell-type specific transcriptome of Npas1+ neurons within the VP. We identified enriched genes, biological processes and transcription factors that may point to a potential novel function of Npas1+ neurons distinct from Npas1- VP neurons. Sox6 being a specific transcription factor is particularly interesting, as a recent anatomical study using Sox6-Cre mice suggests that Sox6+ neurons are a distinct type of neurons within the GPe (Abecassis et al., 2020). Furthermore, this study also revealed that about 93% of Npas1+ neurons within the GP are Sox6+, suggesting a possible functional relationship between these transcription factors. Whether such anatomical description is true for VP Npas1+ neurons remain unknown. Cell-type specific transcriptome analysis also revealed that VP Npas1+ neurons differentially express some genes related to glutamate synthesis and release, while the expression of genes related to GABA neurotransmission are not enriched when compared to total VP. In this context, we highlight the expression of slc17a6 gene, which encodes the transcription of the vesicular glutamate transporter 2 (VGLUT2) (Hayashi et al., 2001; El Mestikawy et al., 2011) and the slc38a2 gene, which encodes the sodium-coupled amino acid transporter 2 (SNAT2) (González-González et al., 2005). VGLUT2 is responsible for packing glutamate into vesicles (Shigeri et al., 2004), while SNAT2 is important for the glutamine uptake, a glutamate precursor (Andersen et al., 2021). Corroborating our results, VGLUT2 positive neurons from the VP have functional projections to the LHb and VTA (Faget et al., 2018; Tooley et al., 2018; Liu et al., 2020), a projection pattern similar to those found in our study for VP Npas1+ neurons.

Our data further demonstrate that chemogenetic activation of VP Npas1+ neurons increased susceptibility to social stress in male mice, while the inactivation of these neurons increased resilience to social stress in both male and female mice. The activation of VP Npas1+ neurons also reduced exploratory behaviors evaluated in the EPM and OF. Overall, the behavioral consequences of manipulating activity in the VP are diverse. Some studies demonstrate that inhibiting the entire VP, by increasing GABAergic tonic inhibition in this region, increases FST immobility and reduces sucrose preference (Skirzewski et al., 2011), while activating orexinergic receptors in GABAergic VP neurons reduces FST immobility time and enhances sucrose preference (Ji et al., 2019). Cell specific manipulation studies demonstrate that silencing VP PV+ neurons in socially-stressed mice decreases social avoidance and the increases in immobility on the tail suspension test (Knowland et al., 2017). Further, inhibition of glutamatergic VP neurons projecting to the LHb blocks social avoidance induced by social stress (Liu et al., 2020). In our study we demonstrate that activation of VP Npas1+ neurons increases susceptibility to a SSDS, whereas their inhibition causes a stress resilient response to a repeated social stressor. Together, such results identify distinct neuronal types within the VP that differentially control behavioral responses to stress, similar to other brain regions (Francis and Lobo, 2017; Fox and Lobo, 2019). These findings extend our knowledge of specific VP neuron subtypes in behavioral response to aversive and rewarding stimuli (Wulff et al., 2019).

We observed a reduction in exploratory behavior in the EPM and the OF when activating VP Npas1+ neurons in our study, implicating alterations in anxiety-like phenotypes. These behaviors are in line with our date demonstrating increased susceptibility to stress in this same condition. In humans, anxiety during young life is correlated to increased risk for depression in adulthood (Kalin, 2017), and the prevalence of anxiety and depression is increased in individuals with a negative coping style to stress (Lew et al., 2019; Xiong et al., 2019). Similar to its role in the behavioral responses to stress, different VP neuronal types have opposite effects on anxiety-like behaviors. Increasing the activity of the whole VP with bicuculine (a GABA_A_ receptor antagonist) enhances exploratory behavior (Reichard et al., 2019). Similar outcomes are observed upon VP GABAergic neuron activation (Li et al., 2020b). Both neurotensin and substance P, when administered in the VP, decrease anxiety-like behaviors (Nikolaus et al., 2000; Ollmann et al., 2015), while increasing in VP activity (Napier et al., 1995; Michaud et al., 2000).

Among the brain targets of VP Npas1+ neurons found in our study, the NAc, the VTA, and the LHb are strongly implicated in the development of depression (Knowland and Lim, 2018; Fox and Lobo, 2019). Furthermore, recent results demonstrate that arkypallidal VP neurons (which resemble arkypallidal neurons within the GP that are Npas1+) project to the NAc, releasing GABA onto medium spiny neurons (MSNs) to promote reward consumption and amplify hedonic actions to reward (Vachez et al., 2021). Our studies indicate that global activation of VP Npas1+ neurons has no effect on hedonic behavior as sucrose preference and intake was unchanged in this group. However, investigation into VP Npas1+ neuron inputs to NAc and the role of this population in stress response, reward consumption, and hedonic behavior is warranted given the unique properties of arkypallidal neurons. Further, while evidence suggests that VP equally projects to both MSN subtypes, those enriched in dopamine receptor 1 vs 2, in NAc (Gangarossa et al., 2013; Li et al., 2018) it is unclear if VP Npas1+ neurons project equally to these two MSN subtypes. Given the dichotomous role of these NAc neuron subtypes in social defeat stress outcomes (Francis et al., 2015), further exploration of Npas1+ VP neuron input to MSNs may shed light into stress mediated- mechanisms in these downstream neurons.

VP projections to the LHb are both excitatory and inhibitory (Wulff et al., 2019). Evidence shows that optogenetic activation of glutamatergic VP neurons induces place aversion (Tooley et al., 2018), and increases behavioral despair in response to SDS (Knowland et al., 2017). Considering the possibility that VP Npas1+ neurons release glutamate within the LHb, another hypothesis is that chemogenetic activation of VP Npas1+ neurons is inducing susceptibility to SDS through increasing LHb activity. This is consistent with an increased glutamate marker in the VP Npas1+ neurons. Future studies are needed to confirm the VP Npas1+ projection neurons that are glutamatergic or if these neurons primarily release GABA, similar to these neurons in the GP (Hernandez et al., 2015; Glajch et al., 2016; Abrahao and Lovinger, 2018).

VTA activity is altered in subjects susceptible to the SDS (Krishnan et al., 2007; Cao et al., 2010; Chaudhury et al., 2013). Following social stress, these individuals have increased VTA dopaminergic neuron firing, specifically to the NAc (Chaudhury et al., 2013). VP activity to VTA is also altered after stress exposure. VP projections to VTA are both glutamatergic and GABAergic and target both dopaminergic and GABAergic interneurons in the VTA, respectively. Increased activity of VP projecting neurons to VTA increases the activity of dopaminergic VTA neurons and increase the susceptibility to SDS (Knowland et al., 2017). It is plausible that VP Npas1+ projections to the VTA increase the activity of dopaminergic neurons through glutamate release, and thus increase susceptibility to SDS.

Our transcriptome data may also provide insight into potential molecular mechanisms involved in VP Npas1+ neuron stress response. Smad2 is a transcriptional modulator upstream to many genes that we found to be enriched in VP Npas1+ neurons (top GO terms, Neurogenesis and Development of Nervous System). Interestingly, Smads (including Smad2) are the main signal transductors activated by the TGF-beta signaling (Hiew et al., 2021), which have being implicated in the effects of stress and depression. For example, TGF-beta mRNA expression is reduced in the amygdala of Wistar rats exposed to chronic mild stress (Bialek et al., 2021). Further, chronic treatment of clinically used antidepressants in rodents increases TGF-beta signaling in the frontal cortex and hippocampus (Dow, 2005; Trojan et al., 2017). Additionally, TGF-beta-Smad signaling is involved in the long term cellular plasticity in reward circuitry that drives drug seeking behavior (Gancarz et al., 2015). The effects of (R)-ketamine treatment on reducing behavioral alterations induced by social defeat stress are blocked when TGF-beta signaling is inhibited (Zhang et al., 2020). Since Npas1+ neurons have an enrichment in genes related to the TGF-Beta/Smad signaling, such mechanisms in these neurons may underlie effects of stress response.

We found some genes (Arhgef10 and Arghgef11) that encodes the Rho guanine nucleotide exchange factors 10 and 11, key regulators of the GTPase RhoA activity (Heasman and Ridley, 2008; Cook et al., 2014), to be enriched in VP Npas1+ neurons. RhoA and its effectors are crucial regulators of dendritic structure and therefore, neuronal plasticity (Nakayama et al., 2000; Newey et al., 2005; Chen and Firestein, 2007). Dendritic atrophy and alterations in dendritic structural plasticity occur in many brain regions after stress exposure in rodents and depression in humans (Anacker et al., 2016; Belleau et al., 2019). Further, our group demonstrated that RhoA play a role in SDS-induced dendritic atrophy in D1-MSNs (Francis et al., 2019; Fox et al., 2020a). Chronic hypoxia exposure increases FST immobility and is related to upregulation of RhoA signaling pathway (Li et al., 2020a), while the inhibition of Rho-associated kinases reverses corticosterone-induced increase in FST immobility and sucrose preference decrease (Wróbel et al., 2018). In rats, the chronic unpredictable stress induces an increase in RhoA levels in the hippocampus (Zhu et al., 2018). Those enriched genes related to the regulation of RhoA activity could also be involved in the mechanisms underlying stress responses in the Npas1+ neurons.

## Conclusions

In conclusion, we have characterized the molecular and circuity projection pattern of VP Npas1+ neurons. We further show that bidirectional chemogenetic modulation of Npas1+ neurons in the VP modulates the outcomes to social defeat stress. The VP has an established role in stress response, as do many of the downstream projection targets. Further, enriched molecules in VP Npas1+ neurons are implicated in stress response and depression in humans. Future studies can elucidate the VP Npas1+ neuron circuitry and molecular mechanisms responsible for these stress outcomes.

## Acknowledgments and Disclosures

This work was supported by NIH R01MH106500, R01DA038613, and R01DA047843 to MKL; R01NS069777 and R01MH112768 to CSC; grants no. 2015/25308-3 and 2018/05496-8 from São Paulo Research Foundation (FAPESP) to GM-S and financed in part by the Coordenação de Aperfeiçoamento de Pessoal de Nível Superior - Brasil (CAPES) - Finance Code 001. FAPESP and CAPES had no further role in the study design; in the collection, analysis, or interpretation of data; in the writing of the report; or in the decision to submit the paper for publication.

## References

Abecassis ZA, Berceau BL, Win PH, García D, Xenias HS, Cui Q, Pamukcu A, Cherian S, Hernández VM, Chon U, Lim BK, Kim Y, Justice NJ, Awatramani R, Hooks BM, Gerfen CR, Boca SM, Chan CS (2020) Npas1 + -Nkx2.1 + Neurons Are an Integral Part of the Cortico-pallido-cortical Loop. J Neurosci 40:743–768 Available at: http://www.jneurosci.org/lookup/doi/10.1523/JNEUROSCI.1199-19.2019.

Abrahao KP, Lovinger DM (2018) Classification of GABAergic neuron subtypes from the globus pallidus using wild-type and transgenic mice. J Physiol 596:4219–4235.

Anacker C, Scholz J, O’Donnell KJ, Allemang-Grand R, Diorio J, Bagot RC, Nestler EJ, Hen R, Lerch JP, Meaney MJ (2016) Neuroanatomic Differences Associated with Stress Susceptibility and Resilience. Biol Psychiatry 79:840–849 Available at: http://dx.doi.org/10.1016/j.biopsych.2015.08.009.

Anders S, Huber W (2010) Differential expression analysis for sequence count data. Genome Biol 11:R106 Available at: https://linkinghub.elsevier.com/retrieve/pii/S0021925819672959.

Andersen J V., Markussen KH, Jakobsen E, Schousboe A, Waagepetersen HS, Rosenberg PA, Aldana BI (2021) Glutamate metabolism and recycling at the excitatory synapse in health and neurodegeneration. Neuropharmacology 196:108719 Available at: https://doi.org/10.1016/j.neuropharm.2021.108719.

Belleau EL, Treadway MT, Pizzagalli DA (2019) The Impact of Stress and Major Depressive Disorder on Hippocampal and Medial Prefrontal Cortex Morphology. Biol Psychiatry 85:443–453 Available at: https://doi.org/10.1016/j.biopsych.2018.09.031.

Bialek K, Czarny P, Wigner P, Synowiec E, Barszczewska G, Bijak M, Szemraj J, Niemczyk M, Tota-Glowczyk K, Papp M, Sliwinski T (2021) Chronic mild stress and venlafaxine treatmentwere associated with altered expression level and methylation status of new candidate inflammatory genes in PBMCs and brain structures of wistar rats. Genes (Basel) 12.

Breton JM, Charbit AR, Snyder BJ, Fong PTK, Dias E V., Himmels P, Lock H, Margolis EB (2019) Relative contributions and mapping of ventral tegmental area dopamine and GABA neurons by projection target in the rat. J Comp Neurol 527:916–941 Available at: https://onlinelibrary.wiley.com/doi/abs/10.1002/cne.24572.

Cao J-L, Cooper DC, Han M-H, Covington III HE, Wilkinson MB, Nestler EJ, Friedman AK, Walsh JJ (2010) Mesolimbic dopamine neurons in the brain reward circuit mediate susceptibility to social defeat and antidepressant action. J Neurosci 30:16453–16458.

Chandra R, Calarco CA, Lobo MK (2019) Differential mitochondrial morphology in ventral striatal projection neuron subtypes. J Neurosci Res 97:1579–1589.

Chandra R, Francis TC, Konkalmatt P, Amgalan A, Gancarz AM, Dietz DM, Lobo MK (2015) Opposing Role for Egr3 in Nucleus Accumbens Cell Subtypes in Cocaine Action. J Neurosci 35:7927–7937 Available at: https://www.jneurosci.org/lookup/doi/10.1523/JNEUROSCI.0548-15.2015.

Chandra R, Francis TC, Nam H, Riggs LM, Engeln M, Rudzinskas S, Konkalmatt P, Russo SJ, Turecki G, Iniguez SD, Lobo MK (2017) Reduced Slc6a15 in Nucleus Accumbens D2-Neurons Underlies Stress Susceptibility. J Neurosci 37:6527–6538 Available at: http://www.jneurosci.org/lookup/doi/10.1523/JNEUROSCI.3250-16.2017.

Chaudhury D et al. (2013) Rapid regulation of depression-related behaviours by control of midbrain dopamine neurons. Nature 493:532–536 Available at: http://dx.doi.org/10.1038/nature11713.

Chen H, Firestein BL (2007) RhoA regulates dendrite branching in hippocampal neurons by decreasing cypin protein levels. J Neurosci 27:8378–8386.

Churchill L, Kalivas PW (1994) A topographically organized gamma-aminobutyric acid projection from the ventral pallidum to the nucleus accumbens in the rat. J Comp Neurol 345:579–595.

Cook DR, Rossman KL, Der CJ (2014) Rho guanine nucleotide exchange factors: Regulators of Rho GTPase activity in development and disease. Oncogene 33:4021–4035.

Dow AL (2005) Regulation of Activin mRNA and Smad2 Phosphorylation by Antidepressant Treatment in the Rat Brain: Effects in Behavioral Models. J Neurosci 25:4908–4916 Available at: https://www.jneurosci.org/lookup/doi/10.1523/JNEUROSCI.5155-04.2005.

El Mestikawy S, Wallén-Mackenzie Å, Fortin GM, Descarries L, Trudeau LE (2011) From glutamate co-release to vesicular synergy: Vesicular glutamate transporters. Nat Rev Neurosci 12:204–216.

Engeln M, Song Y, Chandra R, La A, Fox ME, Evans B, Turner MD, Thomas S, Francis TC, Hertzano R, Lobo MK (2020) Individual differences in stereotypy and neuron subtype translatome with TrkB deletion. Mol Psychiatry Available at: http://dx.doi.org/10.1038/s41380-020-0746-0.

Faget L, Zell V, Souter E, McPherson A, Ressler R, Gutierrez-Reed N, Yoo JH, Dulcis D, Hnasko TS (2018) Opponent control of behavioral reinforcement by inhibitory and excitatory projections from the ventral pallidum. Nat Commun 9:849 Available at: http://dx.doi.org/10.1038/s41467-018-03125-y.

Fox ME, Chandra R, Menken MS, Larkin EJ, Nam H, Engeln M, Francis TC, Lobo MK (2020a) Dendritic remodeling of D1 neurons by RhoA/Rho-kinase mediates depression-like behavior. Mol Psychiatry 25:1022–1034 Available at: http://www.nature.com/articles/s41380-018-0211-5.

Fox ME, Figueiredo A, Menken MS, Lobo MK (2020b) Dendritic spine density is increased on nucleus accumbens D2 neurons after chronic social defeat. Sci Rep 10:12393 Available at: https://doi.org/10.1038/s41598-020-69339-7.

Fox ME, Lobo MK (2019) The molecular and cellular mechanisms of depression: a focus on reward circuitry. Mol Psychiatry Available at: http://www.nature.com/articles/s41380-019-0415-3.

Francis TC, Chandra R, Friend DM, Finkel E, Dayrit G, Miranda J, Brooks JM, Iñiguez SD, O’Donnell P, Kravitz A, Lobo MK (2015) Nucleus accumbens medium spiny neuron subtypes mediate depression-related outcomes to social defeat stress. Biol Psychiatry 77:212–222.

Francis TC, Gaynor A, Chandra R, Fox ME, Lobo MK (2019) The Selective RhoA Inhibitor Rhosin Promotes Stress Resiliency Through Enhancing D1-Medium Spiny Neuron Plasticity and Reducing Hyperexcitability. Biol Psychiatry 85:1001–1010 Available at: https://doi.org/10.1016/j.biopsych.2019.02.007.

Francis TC, Lobo MK (2017) Emerging Role for Nucleus Accumbens Medium Spiny Neuron Subtypes in Depression. Biol Psychiatry 81:645–653 Available at: http://dx.doi.org/10.1016/j.biopsych.2016.09.007.

Gancarz AM, Wang ZJ, Schroeder GL, Damez-Werno D, Braunscheidel KM, Mueller LE, Humby MS, Caccamise A, Martin JA, Dietz KC, Neve RL, Dietz DM (2015) Activin receptor signaling regulates cocaine-primed behavioral and morphological plasticity. Nat Neurosci 18:959–961.

Gangarossa G, Espallergues J, de Kerchove d’Exaerde A, El Mestikawy S, Gerfen CR, Hervé D, Girault J-A, Valjent E (2013) Distribution and compartmental organization of GABAergic medium-sized spiny neurons in the mouse nucleus accumbens. Front Neural Circuits 7:1–20 Available at: http://journal.frontiersin.org/article/10.3389/fncir.2013.00022/abstract.

Glajch KE, Kelver DA, Hegeman DJ, Cui Q, Xenias HS, Augustine EC, Hernandez VM, Verma N, Huang TY, Luo M, Justice NJ, Chan CS (2016) Npas1+ Pallidal Neurons Target Striatal Projection Neurons. J Neurosci 36:5472–5488 Available at: http://www.jneurosci.org/cgi/doi/10.1523/JNEUROSCI.1720-15.2016.

Golden SA, Covington HE, Berton O, Russo SJ (2011) A standardized protocol for repeated social defeat stress in mice. Nat Protoc 6:1183–1191 Available at: http://www.nature.com/doifinder/10.1038/nprot.2011.361.

González-González IM, Cubelos B, Giménez C, Zafra F (2005) Immunohistochemical localization of the amino acid transporter SNAT2 in the rat brain. Neuroscience 130:61–73 Available at: https://linkinghub.elsevier.com/retrieve/pii/S0306452204008152.

Hayashi M, Otsuka M, Morimoto R, Hirota S, Yatsushiro S, Takeda J, Yamamoto A, Moriyama Y (2001) Differentiation-associated Na+-dependent Inorganic Phosphate Cotransporter (DNPI) Is a Vesicular Glutamate Transporter in Endocrine Glutamatergic Systems. J Biol Chem 276:43400–43406 Available at: http://dx.doi.org/10.1074/jbc.M106244200.

Heasman SJ, Ridley AJ (2008) Mammalian Rho GTPases: New insights into their functions from in vivo studies. Nat Rev Mol Cell Biol 9:690–701.

Hernandez VM, Hegeman DJ, Cui Q, Kelver DA, Fiske MP, Glajch KE, Pitt JE, Huang TY, Justice NJ, Chan CS (2015) Parvalbumin+ Neurons and Npas1+ Neurons Are Distinct Neuron Classes in the Mouse External Globus Pallidus. J Neurosci 35:11830–11847 Available at: http://www.jneurosci.org/cgi/doi/10.1523/JNEUROSCI.4672-14.2015.

Hiew LF, Poon CH, You HZ, Lim LW (2021) Tgf-β/smad signalling in neurogenesis: Implications for neuropsychiatric diseases. Cells 10.

Hunt AJ, Dasgupta R, Rajamanickam S, Jiang Z, Beierlein M, Chan CS, Justice NJ (2018) Paraventricular hypothalamic and amygdalar CRF neurons synapse in the external globus pallidus. Brain Struct Funct 223:2685–2698 Available at: http://dx.doi.org/10.1007/s00429-018-1652-y.

Iñiguez SD, Flores-Ramirez FJ, Riggs LM, Alipio JB, Garcia-Carachure I, Hernandez MA, Sanchez DO, Lobo MK, Serrano PA, Braren SH, Castillo SA (2018) Vicarious Social Defeat Stress Induces Depression-Related Outcomes in Female Mice. Biol Psychiatry 83:9–17 Available at: http://dx.doi.org/10.1016/j.biopsych.2017.07.014.

Janky R, Verfaillie A, Imrichová H, Van de Sande B, Standaert L, Christiaens V, Hulselmans G, Herten K, Naval Sanchez M, Potier D, Svetlichnyy D, Kalender Atak Z, Fiers M, Marine J-C, Aerts S (2014) iRegulon: From a Gene List to a Gene Regulatory Network Using Large Motif and Track Collections Bussemaker HJ, ed. PLoS Comput Biol 10:e1003731 Available at: https://dx.plos.org/10.1371/journal.pcbi.1003731.

Ji MJ, Zhang XY, Chen Z, Wang JJ, Zhu JN (2019) Orexin prevents depressive-like behavior by promoting stress resilience. Mol Psychiatry 24:282–293 Available at: http://dx.doi.org/10.1038/s41380-018-0127-0.

Kalin NH (2017) Mechanisms underlying the early risk to develop anxiety and depression: A translational approach. Eur Neuropsychopharmacol 27:543–553 Available at: http://dx.doi.org/10.1016/j.euroneuro.2017.03.004.

Khan HA, Urstadt KR, Mostovoi NA, Berridge KC (2020) Mapping excessive “disgust” in the brain: Ventral pallidum inactivation recruits distributed circuitry to make sweetness “disgusting.” Cogn Affect Behav Neurosci 20:141–159 Available at: http://link.springer.com/10.3758/s13415-019-00758-4.

Kim D, Pertea G, Trapnell C, Pimentel H, Kelley R, Salzberg SL (2013) TopHat2: accurate alignment of transcriptomes in the presence of insertions, deletions and gene fusions. Genome Biol 14:R36 Available at: http://genomebiology.biomedcentral.com/articles/10.1186/gb-2013-14-4-r36.

Klitenick MA, Deutch AY, Churchill L, Kalivas PW (1992) Topography and functional role of dopaminergic projections from the ventral mesencephalic tegmentum to the ventral pallidum. Neuroscience 50:371–386 Available at: https://linkinghub.elsevier.com/retrieve/pii/030645229290430A.

Knowland D, Lilascharoen V, Pacia CP, Shin S, Wang EHJ, Lim BK (2017) Distinct Ventral Pallidal Neural Populations Mediate Separate Symptoms of Depression. Cell 170:284–297.e18 Available at: http://dx.doi.org/10.1016/j.cell.2017.06.015.

Knowland D, Lim BK (2018) Circuit-based frameworks of depressive behaviors: The role of reward circuitry and beyond. Pharmacol Biochem Behav 174:42–52 Available at: https://doi.org/10.1016/j.pbb.2017.12.010.

Krishnan V et al. (2007) Molecular Adaptations Underlying Susceptibility and Resistance to Social Defeat in Brain Reward Regions. Cell 131:391–404 Available at: http://linkinghub.elsevier.com/retrieve/pii/S0092867407012068.

Kuo H, Chang HT (1992) Ventral pallido-striatal pathway in the rat brain: A light and electron microscopic study. J Comp Neurol 321:626–636 Available at: http://doi.wiley.com/10.1002/cne.903210409.

Kupchik YM, Brown RM, Heinsbroek JA, Lobo MK, Schwartz DJ, Kalivas PW (2015) Coding the direct/indirect pathways by D1 and D2 receptors is not valid for accumbens projections. Nat Neurosci 18:1230–1232 Available at: http://dx.doi.org/10.1038/nn.4068.

Lew B, Huen J, Yu P, Yuan L, Wang D-F, Ping F, Abu Talib M, Lester D, Jia C-X (2019) Associations between depression, anxiety, stress, hopelessness, subjective well-being, coping styles and suicide in Chinese university students. PLoS One 14:e0217372.

Li B, Xu Y, Quan Y, Cai Q, Le Y, Ma T, Liu Z, Wu G, Wang F, Bao C, Li H (2020a) Inhibition of RhoA/ROCK pathway in the early stage of hypoxia ameliorates depression in mice via protecting myelin sheath. ACS Chem Neurosci 11:2705– 2716.

Li Y-D, Luo Y-J, Xu W, Ge J, Cherasse Y, Wang Y-Q, Lazarus M, Qu W-M, Huang Z-L (2020b) Ventral pallidal GABAergic neurons control wakefulness associated with motivation through the ventral tegmental pathway. Mol Psychiatry Available at: http://dx.doi.org/10.1038/s41380-020-00906-0.

Li Z, Chen Z, Fan G, Li A, Yuan J, Xu T (2018) Cell-Type-Specific Afferent Innervation of the Nucleus Accumbens Core and Shell. Front Neuroanat 12:1–16 Available at: https://www.frontiersin.org/article/10.3389/fnana.2018.00084/full.

Liu B, Cao Y, Wang J, Dong J (2020) Excitatory transmission from ventral pallidum to lateral habenula mediates depression. World J Biol Psychiatry 2975.

Liu Q-R, Rubio FJ, Bossert JM, Marchant NJ, Fanous S, Hou X, Shaham Y, Hope BT (2014) Detection of molecular alterations in methamphetamine-activated Fos-expressing neurons from a single rat dorsal striatum using fluorescence-activated cell sorting (FACS). J Neurochem 128:173–185 Available at: https://onlinelibrary.wiley.com/doi/10.1111/jnc.12381.

Ma TP, Geyer HL (2018) The Basal Nuclei. In: Fundamental Neuroscience for Basic and Clinical Applications, pp 377–393.e1. Elsevier. Available at: https://linkinghub.elsevier.com/retrieve/pii/B9780323396325000268.

Maere S, Heymans K, Kuiper M (2005) BiNGO: a Cytoscape plugin to assess overrepresentation of Gene Ontology categories in Biological Networks. Bioinformatics 21:3448–3449 Available at: https://academic.oup.com/bioinformatics/article-lookup/doi/10.1093/bioinformatics/bti551.

Mi H, Muruganujan A, Casagrande JT, Thomas PD (2013) Large-scale gene function analysis with the PANTHER classification system. Nat Protoc 8:1551–1566 Available at: http://www.nature.com/articles/nprot.2013.092.

Michaud J-C, Gueudet C, Soubrié P (2000) Effects of neurotensin receptor antagonists on the firing rate of rat ventral pallidum neurons. Neuroreport 11:1437–1441 Available at: http://journals.lww.com/00001756-200005150-00017.

Miller JM, Vorel SR, Tranguch AJ, Kenny ET, Mazzoni P, van Gorp WG, Kleber HD (2006) Anhedonia After a Selective Bilateral Lesion of the Globus Pallidus. Am J Psychiatry 163:786–788 Available at: http://psychiatryonline.org/doi/abs/10.1176/ajp.2006.163.5.786.

Morais-Silva G, Fernandes-Santos J, Moreira-Silva D, Marin MT (2016) Concomitant stress potentiates the preference for, and consumption of, ethanol induced by chronic pre-exposure to ethanol. Brazilian J Med Biol Res 49:1–9 Available at: http://www.ncbi.nlm.nih.gov/pubmed/26628398.

Moussawi K, Kalivas PW, Lee JW (2016) Abstinence From Drug Dependence After Bilateral Globus Pallidus Hypoxic-Ischemic Injury. Biol Psychiatry 80:e79–e80 Available at: http://dx.doi.org/10.1016/j.biopsych.2016.04.005.

Murrough JW, Henry S, Hu J, Gallezot J-D, Planeta-Wilson B, Neumaier JF, Neumeister A (2011) Reduced ventral striatal/ventral pallidal serotonin1B receptor binding potential in major depressive disorder. Psychopharmacology (Berl) 213:547–553 Available at: http://link.springer.com/10.1007/s00213-010-1881-0.

Nakayama AY, Harms MB, Luo L (2000) Small GTPases Rac and Rho in the maintenance of dendritic spines and branches in hippocampal pyramidal neurons. J Neurosci 20:5329–5338.

Nam H, Chandra R, Francis TC, Dias C, Cheer JF, Lobo MK (2019) Reduced nucleus accumbens enkephalins underlie vulnerability to social defeat stress. Neuropsychopharmacology 44:1876–1885 Available at: http://dx.doi.org/10.1038/s41386-019-0422-8.

Napier TC, Mitrovic I, Churchill L, Klitenick MA, Lu X-Y, Kalivas PW (1995) Substance P in the ventral pallidum: Projection from the ventral striatum, and electrophysiological and behavioral cinsequences of pallidal substance P. Neuroscience 69:59–70 Available at: https://linkinghub.elsevier.com/retrieve/pii/0306452295002188.

Newey SE, Velamoor V, Govek EE, Van Aelst L (2005) Rho GTPases, dendritic structure, and mental retardation. J Neurobiol 64:58–74.

Nikolaus S, Huston JP, Hasenöhrl RU (2000) Anxiolytic-like effects in rats produced by ventral pallidal injection of both N- and C-terminal fragments of substance P. Neurosci Lett 283:37–40.

Ollmann T, Péczely L, László K, Kovács A, Gálosi R, Kertes E, Kállai V, Zagorácz O, Karádi Z, Lénárd L (2015) Anxiolytic effect of neurotensin microinjection into the ventral pallidum. Behav Brain Res 294:208–214 Available at: http://dx.doi.org/10.1016/j.bbr.2015.08.010.

Onyewuenyi IC, Muldoon MF, Christie IC, Erickson KI, Gianaros PJ (2014) Basal ganglia morphology links the metabolic syndrome and depressive symptoms. Physiol Behav 123:214–222 Available at: http://dx.doi.org/10.1016/j.physbeh.2013.09.014.

Reichard RA, Parsley KP, Subramanian S, Stevenson HS, Schwartz ZM, Sura T, Zahm DS (2019) The lateral preoptic area and ventral pallidum embolden behavior. Brain Struct Funct 224:1245–1265 Available at: http://link.springer.com/10.1007/s00429-018-01826-0.

Root DH, Melendez RI, Zaborszky L, Napier TC (2015) The ventral pallidum: Subregion-specific functional anatomy and roles in motivated behaviors. Prog Neurobiol 130:29–70 Available at: https://linkinghub.elsevier.com/retrieve/pii/S0301008215000271.

Sanz E, Yang L, Su T, Morris DR, McKnight GS, Amieux PS (2009) Cell-type-specific isolation of ribosome-associated mRNA from complex tissues. Proc Natl Acad Sci 106:13939–13944 Available at: http://www.pnas.org/cgi/doi/10.1073/pnas.0907143106.

Shannon P, Markiel A, Ozier O, Baliga NS, Wang JT, Ramage D, Amin N, Schwikowski B, Ideker T (2003) Cytoscape: a software environment for integrated models of biomolecular interaction networks. Genome Res 13:2498–2504 Available at: http://ci.nii.ac.jp/naid/110001910481/.

Shigeri Y, Seal RP, Shimamoto K (2004) Molecular pharmacology of glutamate transporters, EAATs and VGLUTs. Brain Res Rev 45:250–265.

Skirzewski M, López W, Mosquera E, Betancourt L, Catlow B, Chiurillo M, Loureiro N, Hernández L, Rada P (2011) Enhanced GABAergic tone in the ventral pallidum: Memory of unpleasant experiences? Neuroscience 196:131–146 Available at: http://dx.doi.org/10.1016/j.neuroscience.2011.08.058.

Stanco A, Pla R, Vogt D, Chen Y, Mandal S, Walker J, Hunt RF, Lindtner S, Erdman CA, Pieper AA, Hamilton SP, Xu D, Baraban SC, Rubenstein JLR (2014) NPAS1 Represses the Generation of Specific Subtypes of Cortical Interneurons. Neuron 84:940–953 Available at: https://linkinghub.elsevier.com/retrieve/pii/S089662731400960X.

Stout KA, Dunn AR, Lohr KM, Alter SP, Cliburn RA, Guillot TS, Miller GW (2016) Selective Enhancement of Dopamine Release in the Ventral Pallidum of Methamphetamine-Sensitized Mice. ACS Chem Neurosci 7:1364–1373 Available at: https://pubs.acs.org/doi/10.1021/acschemneuro.6b00131.

Stuke H, Hanken K, Hirsch J, Klein J, Wittig F, Kastrup A, Hildebrandt H (2016) Cross-Sectional and Longitudinal Relationships between Depressive Symptoms and Brain Atrophy in MS Patients. Front Hum Neurosci 10:1–8 Available at: http://journal.frontiersin.org/article/10.3389/fnhum.2016.00622/full.

Taylor SR, Badurek S, Dileone RJ, Nashmi R, Minichiello L, Picciotto MR (2014) GABAergic and glutamatergic efferents of the mouse ventral tegmental area. J Comp Neurol 522:3308–3334.

Teh CHL, Lam KKY, Loh CC, Loo JM, Yan T, Lim TM (2006) Neuronal PAS Domain Protein 1 Is a Transcriptional Repressor and Requires Arylhydrocarbon Nuclear Translocator for Its Nuclear Localization. J Biol Chem 281:34617–34629 Available at: http://www.jbc.org/lookup/doi/10.1074/jbc.M604409200.

Tooley J, Marconi L, Alipio JB, Matikainen-Ankney B, Georgiou P, Kravitz A V., Creed MC (2018) Glutamatergic Ventral Pallidal Neurons Modulate Activity of the Habenula–Tegmental Circuitry and Constrain Reward Seeking. Biol Psychiatry 83:1012–1023 Available at: https://doi.org/10.1016/j.biopsych.2018.01.003.

Trojan E, Ślusarczyk J, Chamera K, Kotarska K, Glombik K, Kubera M, Basta-Kaim A (2017) The modulatory properties of chronic antidepressant drugs treatment on the brain chemokine - Chemokine receptor network: A molecular study in an animal model of depression. Front Pharmacol 8:1–16.

Vachez YM, Tooley JR, Abiraman K, Matikainen-Ankney B, Casey E, Earnest T, Ramos LM, Silberberg H, Godynyuk E, Uddin O, Marconi L, Le Pichon CE, Creed MC (2021) Ventral arkypallidal neurons inhibit accumbal firing to promote reward consumption. Nat Neurosci Available at: 10.1038/s41593-020-00772-7.

Wróbel A, Serefko A, Rechberger E, Banczerowska-Górska M, Poleszak E, Dudka J, Skorupska K, Miotła P, Semczuk A, Kulik-Rechberger B, Mandziuk S, Rechberger T (2018) Inhibition of Rho kinase by GSK 269962 reverses both corticosterone-induced detrusor overactivity and depression-like behaviour in rats. Eur J Pharmacol 837:127–136 Available at: https://doi.org/10.1016/j.ejphar.2018.08.027.

Wulff AB, Tooley J, Marconi LJ, Creed MC (2019) Ventral pallidal modulation of aversion processing. Brain Res 1713:62–69 Available at: https://doi.org/10.1016/j.brainres.2018.10.010.

Xiong W, Liu H, Gong P, Wang Q, Ren Z, He M, Zhou G, Ma J, Guo X, Fan X, Liu M, Yang X, Shen Y, Zhang X (2019) Relationships of coping styles and sleep quality with anxiety symptoms among Chinese adolescents: A cross-sectional study. J Affect Disord 257:108–115.

Zhang K, Yang C, Chang L, Sakamoto A, Suzuki T, Fujita Y, Qu Y, Wang S, Pu Y, Tan Y, Wang X, Ishima T, Shirayama Y, Hatano M, Tanaka KF, Hashimoto K (2020) Essential role of microglial transforming growth factor-β1 in antidepressant actions of (R)-ketamine and the novel antidepressant TGF-β1. Transl Psychiatry 10 Available at: http://dx.doi.org/10.1038/s41398-020-0733-x.

Zhu X-L, Chen J-J, Han F, Pan C, Zhuang T-T, Cai Y-F, Lu Y-P (2018) Novel antidepressant effects of Paeonol alleviate neuronal injury with concomitant alterations in BDNF, Rac1 and RhoA levels in chronic unpredictable mild stress rats. Psychopharmacology (Berl) 235:2177–2191 Available at: http://link.springer.com/10.1007/s00213-018-4915-7.

